# Germline-encoded TCR-MHC contacts promote TCR V gene bias in umbilical cord blood T cell repertoire

**DOI:** 10.1101/621821

**Authors:** Kai Gao, Lingyan Chen, Yuanwei Zhang, Yi Zhao, Ziyun Wan, Jinghua Wu, Liya Lin, Yashu Kuang, Jinhua Lu, Xiuqing Zhang, Lei Tian, Xiao Liu, Xiu Qiu

## Abstract

T cells recognize antigens as peptides bound to major histocompatibility complex (MHC) proteins through T cell receptors (TCRs) on their surface. To recognize a wide range of pathogens, each individual possesses a substantial number of TCRs with an extremely high degree of variability. It remains controversial whether germline-encoded TCR repertoire is shaped by MHC polymorphism and, if so, what is the preference between MHC genetic variants and TCR V gene compatibility. To investigate the “net” genetic association between MHC variations and TRBV genes, we applied quantitative trait locus (QTL) mapping to test the associations between MHC polymorphism and TCR β chain V (TRBV) genes usage using umbilical cord blood (UCB) samples of 201 Chinese newborns. We found TRBV gene and MHC loci that are predisposed to interact with one another differ from previous conclusions. The majority of MHC amino acid residues associated with the TRBV gene usage show spatial proximities in known structures of TCR-pMHC complexes. These results show for the first time that MHC variants bias TRBV gene usage in UCB of Chinese ancestry and indicate that germline-encoded contacts influence TCR-MHC interactions in intact T cell repertoires.

## Introduction

T cell immune surveillance is critical for the health of all jawed vertebrates. Most T cells express αβ TCRs and recognize peptides derived from digested proteins when presented at the cell surface in MHC molecules. How αβ TCR interacts with peptide-MHC (pMHC) has been a particularly attractive field as it may contribute to developing strategies for manipulating T cell responses in many diseases including immunodeficiencies, tumor, autoimmune, and allergic diseases.

Looking at the structures, TCR-pMHC interaction is a delicate process. TCR exists in heterodimers and the binding site of each TCR chain can be divided into three complementarity-determining regions (CDRs), called CDR1, 2, and 3. The most variable region CDR3 is formed by somatic recombination of the variable (V), joining (J), and in β chains diversity (D) genes. Less viable regions CDR1 and CDR2 loop sequences are constant for each type of chain, and are therefore referred to as “germline-derived”. Unlike CDR3, CDR1 and CDR2 regions are encoded by only TCR V genes. As such, the T cell repertoire of each individual may have a potential number of 10^18^ TCRs^1^. Human MHC genes are also known as human leukocyte antigen (HLA) genes. HLA genes are extremely polymorphic with more than 12000 known alleles and MHC haplotypes are highly variable in different ethnic groups^2,3^. Each individual inherits one set of MHC genes from a parent including classical MHC class I loci (HLA-A, -B, and -C) and classical MHC class II loci (HLA-DP, -DQ, and -DR). In the dozens of structures of TCR-pMHC complexes that have been solved, CDR1 and CDR2 loops are shown to in contact with the conserved α-helical residues of the MHC molecules, and the highly variable CDR3 loops primarily interact with the peptide^4-6^.

Recently, biased αβTCR repertoires and TCR ‘signatures’ raised against specific antigens have been observed in various diseases, especially in virus infections, autoimmune disorders, and tumor^7^. Little is known about whether, and if so how MHC genotype influences the composition of TCR repertoire. Most recently, a genetic study done by Pritchard laboratory showed that MHC gene is the most influential gene of TCR V gene usage using RNA-seq data from a large cohort of European adults^8^. However, some structural and functional studies proposed that neither MHC residues nor germline-encoded TCR sequences are indispensable for TCR-pMHC interaction. As TCR repertoire can change greatly with T-cell turnover and immune responses to environmental insults, it is critical to determine how MHC molecules affect TCR repertoire with confounding factors in control.

In this study, we attempt to investigate if MHC genetic variations reshape the intact TCR repertoires using umbilical cord blood (UCB) samples from 201 Chinese newborns. UCB samples are used to exclude any influential factors that have yet to be introduced to the TCR repertoires. We sequenced TCR β chain V genes and the MHC region and performed QTL mapping to test the associations between TRBV gene usage and MHC variations at three levels of allele, nucleotide and amino acid. Our results identified TRBV gene and MHC loci that are predisposed to interact with one another. Contacts between MHC and TRBV molecules promote a genotype shift in TRBV gene usage.

## Results

### TRBV gene usage is influenced by MHC genetic variation

To estimate the usage of TRBV genes, we sequenced TCR β chain V genes using DNA collected from UCB of 201 Chinese newborns. Then we calculated the proportion of reads that mapped uniquely to each Vβ gene of all mapped reads. The MHC region of the same individual was sequenced by target capture sequencing and classical four-digit alleles were imputed from SNP data. A schematic overview of the analysis is presented in **Fig. 1A.**

We extracted 48 TRBV genes in each of the 201 samples using immune repertoire sequencing data **(Supplementary Fig. 1, Supplementary table 1 and 2).** Undetected TRBV genes are randomly scattered, which is most likely due to individual differences. Unique alleles of 23 HLA-A, 46 HLA-B, 21 HLA-C, 28 DRB1, 16 DPB1, 4 DPA1, 14 DQB1, and 15 DQA1 are present in our cohort (**Supplementary Table 3).** We then performed QTL mapping between TRBV usage and MHC alleles. As a control, B cell Immunol Globulin Heavy chain V (IGHV)genes, which are not expected to interact with MHC, were analyzed in a similar pipeline. Our results showed that the frequencies of TRBV genes, but not the IGHV genes, are significantly associated with MHC alleles (**Fig.1B, Supplementary Table 4 and 5**). We then performed QTL mapping using single nucleotide polymorphisms (SNPs) (**Supplementary Table 6 and 7**) and amino acid variations (**Supplementary Table 8 and 9)** corresponding to MHC alleles. Again, only TRBV gene usage is significantly associated with MHC at SNP and amino acid levels (**Fig. 1B**). These are the first genetic evidence that TRBV gene usage is significantly influenced by MHC genes in UCB T cell repertoires.

**Figure 1.**
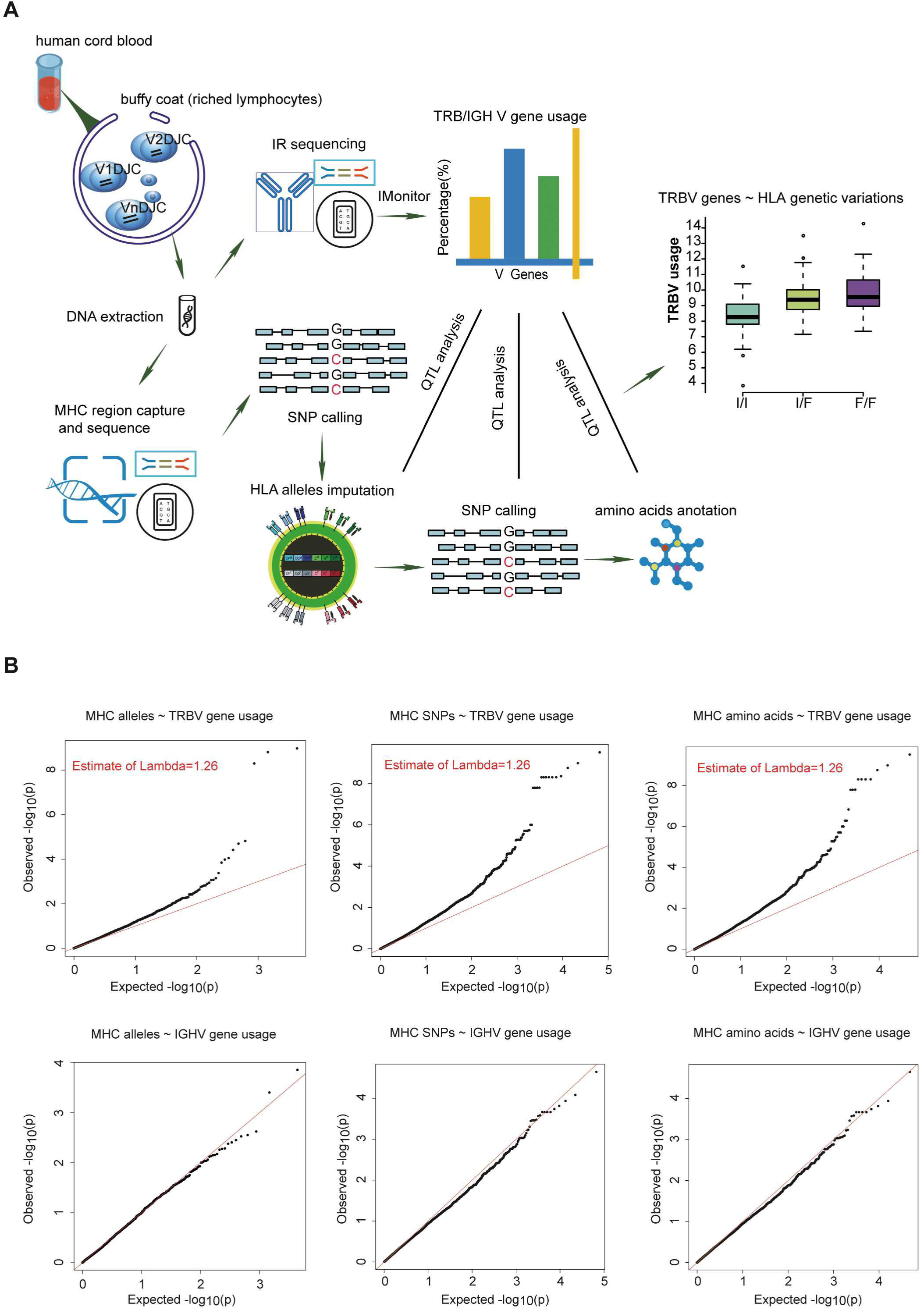
The frequencies of TRBV genes is significantly associated with variations in the MHC locus. **(A)** Schematic overview of the analysis. Usage of TCR β chain V genes was estimated by mapping buffy coat DNA sequencing reads to the human TRBV gene database. MHC alleles were imputed with Beagle. SNPs were called using BWA (version 0.5.9). Amino acid polymorphisms corresponding to MHC alleles were obtained by annotation of SNPs using ANNOVAR. The associations of Vβ usage with nucleotide and amino acid genotypes were tested by QTL mapping using linear regression model. **(B)** QQ plots for the associations between MHC alleles/SNPs/amino acid variations and the usage of TRBV genes (up) or IGHV genes (down). The red line refers to a normal distribution; each black dot represents one allele/SNP/amino acid variation. Estimated inflation factor lambda is provided if it is statistically significant.

### TRBV genes differ in their association with different MHC locus

Next, we aim to find which TRBV genes and MHC variants are most likely to be in strong associations. Our results show that the frequencies of 8.3% (4/48) of TRBV genes can be explained by MHC alleles and the frequencies of 14.6% (7/48) of TRBV genes can be explained by MHC nucleotide and/or amino acid variations (**Fig. 2A**). Among the seven TRBV genes influenced by MHC, TRBV13, TRBV7-6, TRBV7-9, TRBV10-3, and TRBV30 are novel; TRBV20-1 and TRBV9 have been described before in European adults^8^. In accordance with the previous report, our results indicate that TCR V genes differ in their compatibilities with MHC polymorphism.

**Figure 2.**
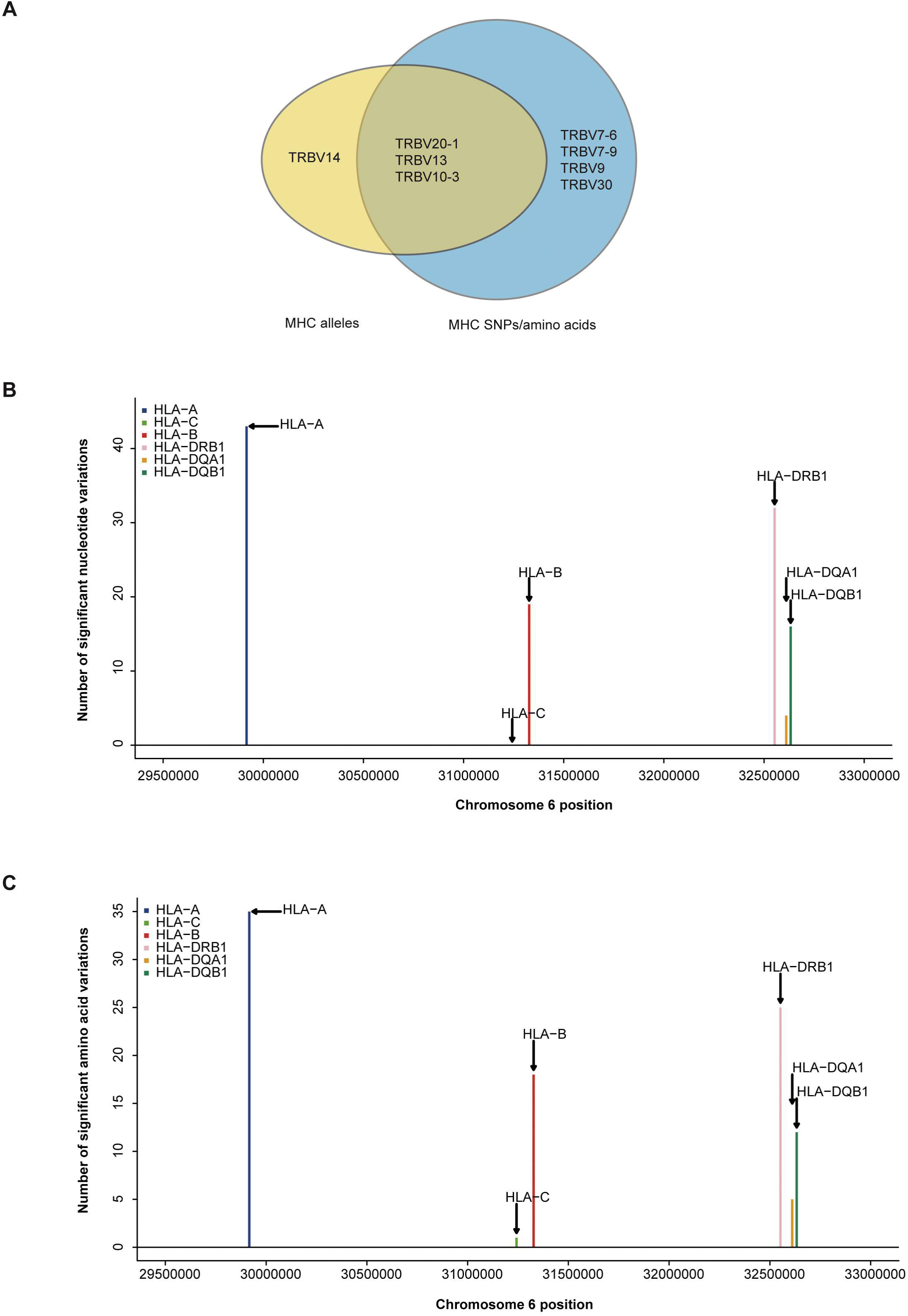
TRBV genes differ in their association with different MHC locus. **(A)** TRBV gene usage explained by MHC alleles (yellow), nucleotide, and amino acid variations of the MHC alleles (blue). The associations of Vβ gene usage with the number of nucleotide variations **(B)**, and with the number of amino acid variations **(C)** (FDR ≤0.05). SNPs are binned according to their genomic position.

Looking at the distribution of significant associations at different MHC loci, 9 MHC alleles, 114 MHC SNPs and 96 MHC amino acid variations are associated with TRBV usage at FDR<0.05 (Supplementary Table 4, 6, and 8). MHC class I locus HLA-A and MHC class II locus DRB1 each has a larger number of variations associated with TRBV genes than that of any other loci of the same class at both nucleotide and amino acid level (**Fig. 2B and 2C**). If the specificity of TCRs towards MHC molecules depends on only the conservative regions of MHC molecules, then we may expect to find the number of TCR associated MHC molecules to be approximately proportional to the number of MHC variations in each region. Interestingly, HLA-C locus has the minimum number of variations in association with TRBV genes despite the fact that the degree of HLA-C loci diversity is as high as that of HLA-A loci. Although HLA-C shares sequence homology with HLA-A and HLA-B molecules, it is substantially different from other MHC I molecules in many different ways. One of the main feature distinguishing HLA-C is its low expression, that the number of surface HLA-C proteins are estimated to be only ∼10% of that of HLA-A and HLA-B molecules^9-11^. Thus, it is highly possible TCR usage bias explained by different MHC loci is also shaped by the intrinsic differences in the abundance of surface MHC proteins, suggesting a strong influence from direct physical contacts.

### Independent MHC residues that bias TRBV gene usage

Due to the strong linkage disequilibrium (LD) structure in the MHC region, it is unclear which MHC variant is responsible for a particular association. Therefore, we performed conditional analysis for each TRBV genes to identify their potential independent MHC amino acid variants. For instance, we examined all TRBV13-associated MHC amino acids. Twenty polymorphic amino acids in HLA-A, five in HLA-B and one in HLA-DRB1 (position 71) showed significant association with TRBV13 (FDR<0.05). Except HLA-A position 97 and HLA-DRB1 position 71, other amino acid positions initially have four potential linkage groups **(Fig. 3A).** Our conditional analysis further confirmed that amino acid position 97 of HLA-A have the strongest association with TRBV13, followed by position 71 of HLA-DRB1 (**Fig. 3 B and C**, conditional *P*< 0.05). After conditioning for these two top positions, no other associations reach a significant level.

**Figure 3.**
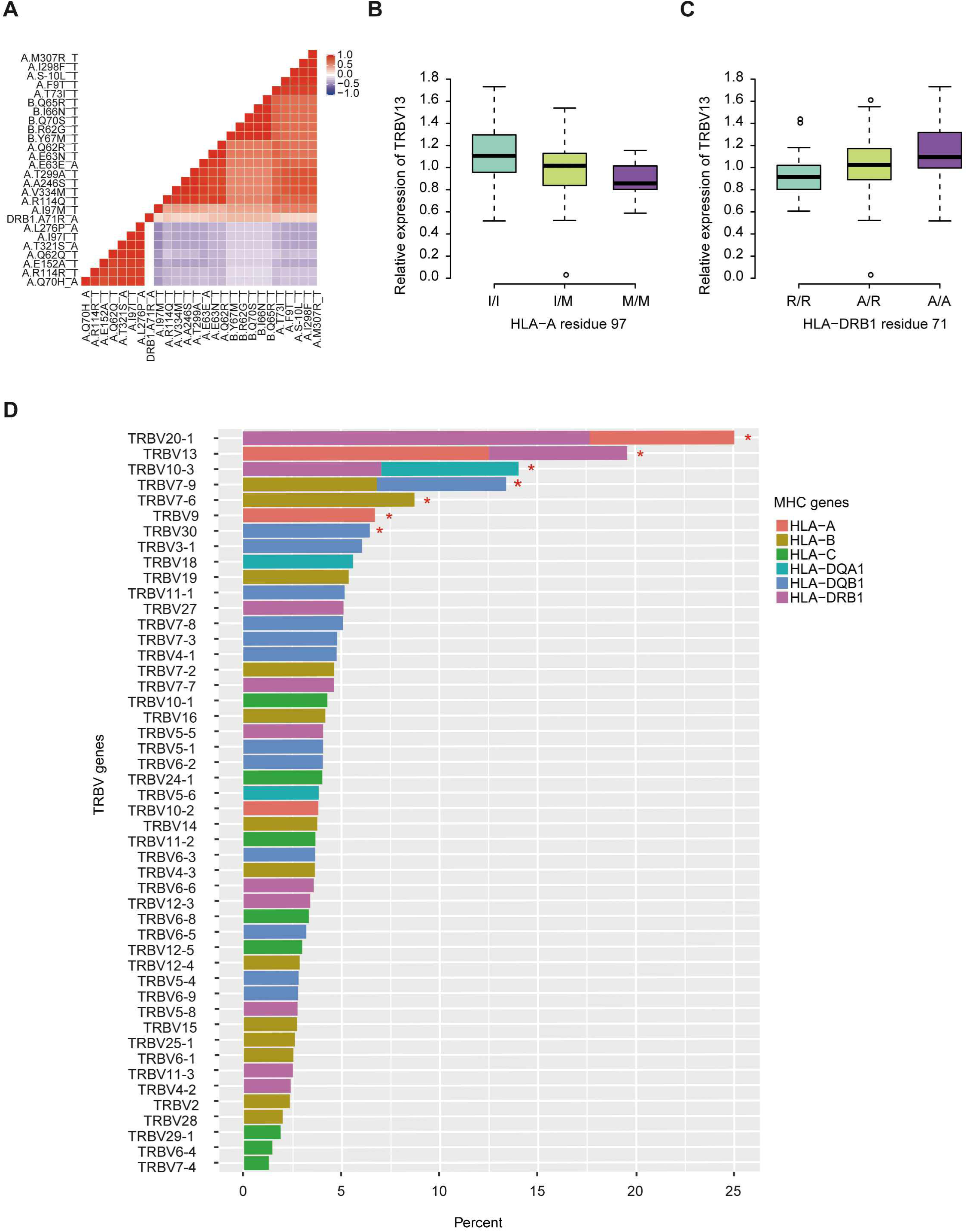
Independent associations between TRBV usage and amino acids variations in MHC alleles. **(A)** The heat map representing a color-coded correlation matrix of all the 26 MHC amino acids that are significantly associated with TRBV13 usage (FDR ≤ 0.05). The frequencies of TRBV13 influence by the two independent amino acid variations at HLA-A residue 97 **(B)** and HLA-DRB1 residue 71 **(C). (D)** Variation of TRBV gene usage explained by MHC amino acids variations of the MHC alleles. Values were adjusted R-square derived from linear regression model. A star indicates that the total proportion of variation explained by the MHC gene components was significant at 5% FDR.

Using this method, 11 independent amino acid variants are found to have the highest possibilities of influencing TRBV gene usage (**Fig. 3, Supplementary Fig. 2**). Three of the 11 independent amino acid variations, HLA-DRB1 residue 67 and 71, HLA-DQB1 residue 55 are associated with type 1 diabetes, multiple sclerosis, hypothyroidism, rheumatoid arthritis, and other inflammatory polyarthritis according to PheWAS database^12, 13^. We then estimated to what extent these MHC variants can influence each TRBV gene usage variation using multiple linear regression. The results showed that 6-23.3% of the proportion of TRBV variability can be explained by MHC amino acid variations, highlighting an important role of MHC in shaping TCR repertoires (**Fig. 3D, Table 1**).

**Table 1.** Usage variation of TRBV genes explained by independent amino acid variations in the MHC locus.

| TRBV Gene ID | Top Signal | Second Signal | % of Variance explained by top signal | % of Variance explained by second signal | % of Total variance explained |
| --- | --- | --- | --- | --- | --- |
| TRBV20-1 | HLA-DRB1.I67F | HLA-A.D116Y | 17.68110065 | 7.374011699 | 23.30052674 |
| TRBV13 * | HLA-A.I97M | HLA-DRB1.A71R | 12.5520188 | 7.0425016 | 17.07336994 |
| TRBV7-6* | HLA-B.V282I | NA | 8.74398907 | NA | 8.74398907 |
| TRBV7-9* | HLA-B.E45K | HLA-DQB1.L75V | 6.829646821 | 6.589435636 | 12.40109952 |
| TRBV10-3* | HLA-DRB1.D57S | HLA-DQA1.A199T | 7.061965751 | 6.994849504 | 15.62058733 |
| TRBV9 | HLA-A.L276P | NA | 6.729255542 | NA | 6.729255542 |
| TRBV30* | HLA-DQB1.R55P | NA | 6.468371558 | NA | 6.468371558 |

\*novel TRBV genes associated with MHC amino acid variations

### TRBV gene associated MHC amino acids locate in TCR binding pockets

MHC residues near the contact interface between TCR and MHC or located in the polymorphic “pockets” of the peptide-binding grooves are most likely to influence the TCR-pMHC (peptide-major histocompatibility complex) interaction^6^. To see whether TRBV gene associated MHC residues are adjacent to or have direct contacts with TCRs on the molecular structure, we mapped these MHC residues onto protein structures of pMHC complex downloaded from PDB database (**Fig. 4A)**. We then collected all structural information of TCR-pMHC complex consisting of 69 HLA-A, 23 HLA-B, 12 HLA-DRB1, 8 HLA-DQA1 or 8 HLA-DQB1 and their paired TCR and peptides (**Supplementary Table 10**). The residues with a high possibility of influencing TRBV gene usage tend to be either physically near or in direct contact with the TCR in structures (**Fig. 4B**). For instance, HLA-DRB1 residue at position 67 predicted in our model with the highest association with TRBV20-1 showed contact with TCR in 42% analyzed complex and with peptide in 75% analyzed complex. These results suggest that MHC amino acid residues influence TCR-pMHC interactions via direct physical contacts.

**Figure 4.**
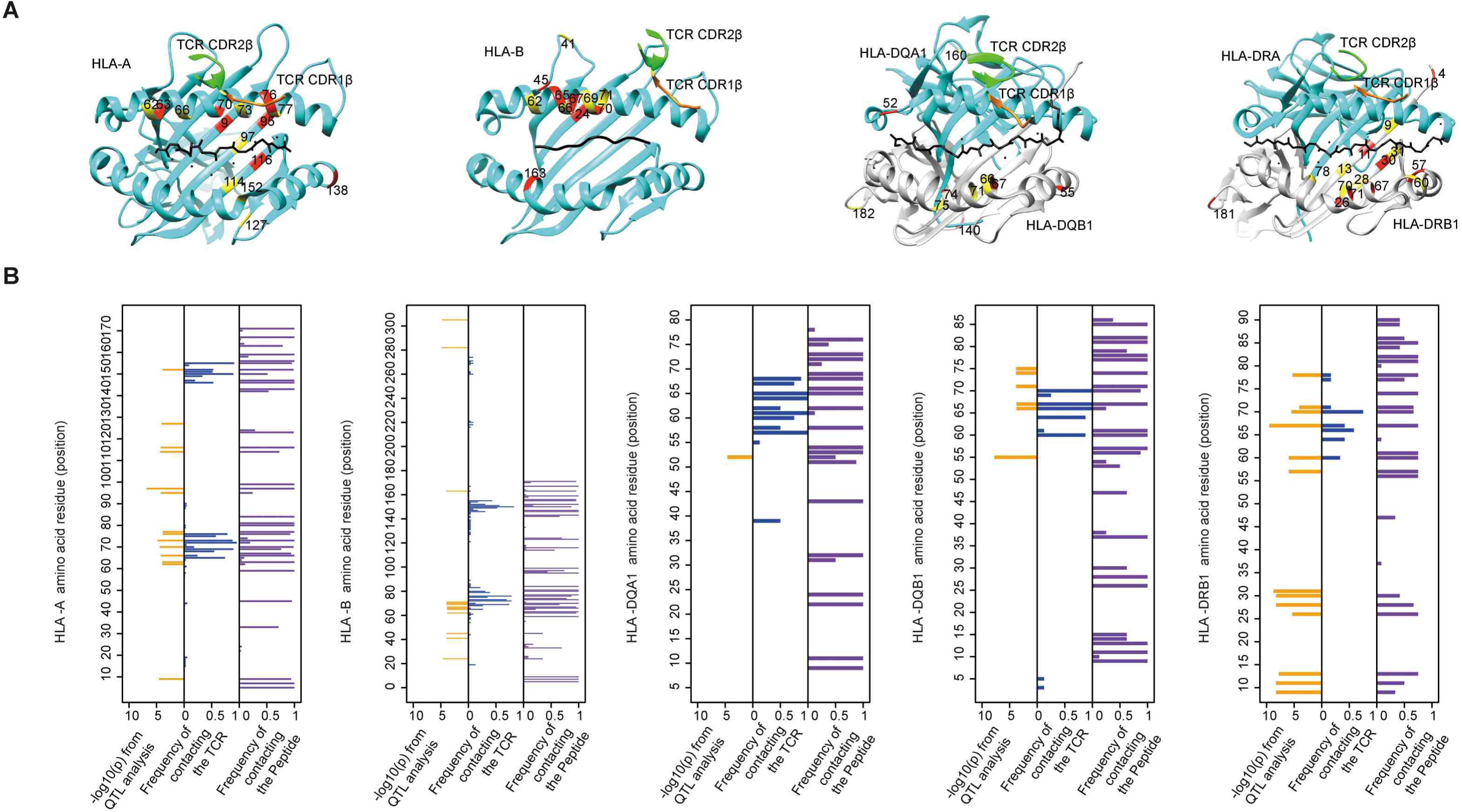
The associated MHC residues tend to be at the TCR-pMHC interface. **(A)** Mapping MHC amino acids biasing the TRBV genes usage (red and yellow) onto a structure of MHC genes (cyan) with CDR1 β chain (orange) and CDR2 β chain (green) (PDB ID from left to right: 2BNQ, 4G9F, 4OZF, and 1J8H). **(B)** Comparison of the Log10-transformed P values for the QTL analysis (left in each panel) and the frequency with which these amino acids physically contact the TCR (middle in each panel) or the Peptide (right in each panel) in solved complexes (from left to right: 69 HLA-A complexes, 23 HLA-B complexes, 8 HLA-DQA1 complexes, 8 HLA-DQB1 complexes, and 12 HLA-DRB1 complexes).

## Discussion

Our results show that MHC polymorphism plays an important role in shaping UCB TCR repertoires. 14.6% (7/48) of TRBV genes are significantly associated with nucleotide and/or amino acid variations of the MHC molecules. Among these TRBV associated MHC loci, we are able to pinpoint 11 independent influential MHC amino acid residues, the majority of which are located in HLA-A and HLA-DRB1 loci. The structural analysis confirmed that the majority of TRBV associated MHC residues are positioned at the TCR-pMHC contact interfaces of known protein complexes, which indicates that MHC molecules have a higher probability of influencing TRBV gene frequencies through physical contacts. In summary, we conclude that MHC variations sculpt UCB TRBV gene repertoires by favoring more compatible TCR-MHC pairs in thymic selection.

Our results shed light on the long debate about the basis of TCR specificity for MHC molecules. In 1971, Jerne proposed that TCR and MHC genes coevolve to have inherent predisposition to interact with one another^14^. Jerne’s idea was further extended that TCRs’ biases for MHC are dictated by conserved CDR1 and CDR2 loops encoded by TCR V genes^15^, which was later validated in many known TCR-pMHC complexes^4,16-19^. In contradiction to this theory, a human TCR complex with an MHC class II molecule demonstrated no dependent of CDR1 and CDR2 for MHC recognition^20^. In this complex, CDR1 and CDR2 have a few contacts with any kind of MHC, and CDR3s have extensive contacts with the peptide and the α helices^20^. It is currently impossible, or probably unnecessary, to thoroughly exam all possible TCR-pMHC structures to conceive the rules for TCR-pMHC interactions. Our results provided genetic evidence that intrinsic TCR-MHC bias exists in the UCB samples of a relatively large cohort of newborns. A recent study consistent with this idea comes from Pritchard laboratory, which utilized RNA-seq data from adult peripheral blood^8^. In summary, our genetic results are supportive of the inherited reactivity of TCR-MHC molecules.

Previous studies on the TCR bias for MHC molecules have utilized adult peripheral blood samples of mouse and human. When using adult samples, precautions need to be taken into consideration to exclude confounding factors that may substantially shape T cell repertoire. Two such factors are the thymic involution during aging and prevalent pathogen infections. To minimize any known and unknown confounding factors, we utilized the newborns’ UCB samples that allow us to assess the “net” genetic association between TCR and MHC. The results of our UCB data differ from previous data: HLA-A and HLA-B, instead of HLA-C genes, are found to be the most influential MHC class I loci to TRBV gene usage. Although the conservative regions of HLA-C proteins are similar to that of HLA-A and HLA-B proteins, HLA-C is substantially different from other MHC I molecules, especially the surface expression of HLA-C proteins is much lower than those of the HLA-A and HLA-B proteins^21^. In addition, only a minority of CD8T cells are restricted by HLA-C, and they have recently been found to be critical in response to chronic infections such as Epstein-Barr virus and HIV infection^22^. Therefore, the associations between HLA-C and TCRs found in the circulation of adults may be biologically meaningful, but may also reflect the prevalence of HLA-C restricted T cell responses.

It is interesting that several MHC loci stand out as more influential than other loci in shaping TRBV genes. There may be important biological meanings in the selective involvement of certain MHC loci in thymic education. The conventional TCR-MHC system is not the only function of the MHC genes or necessarily the original function from an evolution point of view. For instance, many non-classical MHC class I molecules, as well as HLA-C, also function as a ligand for killer immunoglobulin receptors (KIRs) to regulate nature killer (NK) cell activities. Crystal structures of the KIR2DL2-HLA-Cw3 and KIR2DL1-HLA-Cw4 complexes revealed the precise contacting site of HLA-C with KIR molecules, which is predicted to result in KIR competing with TCR for HLA-C interaction^11^. Similarly, in the risk predictions of MHC mismatching transplantations, HLA-A, -B, -C and -DRB1 mismatches are associated with higher risks of graft-versus-host response than those of other classical MHC loci^23^. Together, these results suggest a modification to the applicable MHC loci of the coevolution theory.

With regard to the TRBV genes, we found seven MHC biased TRBV genes among which, TRBV13, TRBV7-6, TRBV7-9, TRBV10-3, and TRBV30 are first reported, and TRBV20-1 and TRBV9 have been described before in European adults^8^. Notably, structures that composed of TRBV13 has been extensively studied in mouse. Complexes that include TRBV13-2 show that amino acids in its CDR2 loop react with related sites on the MHCII α1 helix despite various docking angles of the TCR, and TRBV13-1 CDR1 and CDR2 loops have more than one docking site on α1 helix and shifts according to docking positions^24^. A dynamic interplay between TCR and MHC molecules has gathered more and more evidence. It is possible that we could eventually construct a list of conserved interactions between TCR and MHC with genetic studies, even though, the two molecules may not interact in a conventional way.

In summary, our results showed that TCR Vβ genes are significantly associated with the MHC genotypes in the UCB from a cohort of 201 Chinese newborns. Our structure analysis suggests that MHC amino acid residues associated with the TRBV gene usage are in contact or adjacent with the TCRβ chains in physical structure showing the substantial potential of MHC to influence TCR in protein-protein interaction. Our results confirm and extend our knowledge of TCR-MHC association and contribute to the richness of side-by-side comparisons of population-based studies as a strategy to further understanding the nature of TCR-MHC interactions and its implications in human.

## Methods

### Umbilical cord blood collection and DNA isolation

6ml umbilical cord blood was collected from the unborn placenta of full-term deliveries in each of 201 healthy newborns at Guangzhou Women and Children’s Medical Center (Guangzhou city, Guangdong, China). The sample collection was performed in accordance with the ethical standards of the Ethics Committee of Guangzhou Women and Children’s Medical Center (GWCMC), and written informed consent was obtained from all participating pregnant women. The buffy coat of cord blood was freshly isolated with density gradient centrifugation and DNA was extracted using HiPure Blood DNA Mini Kit (Magen) according to the manufacturer’s protocol. DNA concentration and integrity were measured by Qubit 3.0 fluorometer (Life Technologies, Paisley, UK) and agarose gel (Agilent) electrophoresis.

### Immune repertoire library preparation and sequencing

1.2ug DNA sample was partitioned to construct a library for TCR β (TRB) chain and immune globulin heavy (IGH) chain sequencing using a two-step-PCR method. Firstly, buffy coat DNA was subjected to Multiplex PCR (MPCR) using published primers and cycling conditions ^<u>25</u>^ to enrich rearranged complementarity determining region 3 (CDR3) of the variable regions of TRB/IGH chain. The second PCR introduced the sequencing primers, Illumina primers P5 and P7, into the first PCR products. IGH and TRB genes were sequenced on the HiSeq 4000 (Illumina, La Jolla, CA) with the standard paired-end 150 and paired-end 100 protocol, respectively. Base calling was performed according to the manufacturer’s instruction.

### MHC region capture, library construction and sequencing

MHC region capture sequencing was performed using the similar protocol reported before^26^ with the same design of MHC capture probes named as 110729_HG19_MHC_L2R_D03_EZ, whose product information is provided on the Roche NimbleGen website (https://sequencing.roche.com/en/technology-research/research/immunogenetics.html).

However, sequencing adaptors are modified to fit BGISEQ-500 sequencer (BGI-Shenzhen, Shenzhen, China). In brief, 1ug shotgun library was hybridized to the capture probes following the manufacturer’s protocols (Roche NimbleGen) and the captured target fragments were amplified using AccuPrime® Pfx DNA Polymerase (Invitrogen). The PCR products of captured DNA were purified and quantified by Qubit® dsDNA BR Assay Kits (Invitrogen), and then used to construct sequencing library guided by the manufacturer’s protocol (BGISEQ-500) ^27^ including cyclizing and digestion with enzymes and quantified by Qubit® ssDNA Assay Kit (Invitrogen). Finally, libraries were sequenced with standard paired-end 50 reads on the BGISEQ-500 sequencer following manufacturer’s instructions ^27^.

### Immune repertoire analysis

TRB and IGH Sequencing data were analyzed using IMonitor (version 1.3.0) ^25^, of which the specific parameters are as the followings: -ec -k 100 (IGH pipeline is 150) -jif 80 -vif 80 -v 33 -d. Other parameters are using the defaults. To allow data analysis by QTL mapping, undetectable TRBV and IGHV gene were defined as zero. Log2-transformed usage of TRBV (0.01 pseudo-usage was added to avoid zeroes for TRBV) and IGHV (0.00001 pseudo-usage was added to avoid zeroes for IGHV) were displayed by R code with hierarchical clustering of both rows and columns.

### Variant calling in raw reads of MHC region

MHC targeted capture sequencing data was first quality controlled by filtering out reads with more than 50% bad bases that have quality score <=5, then aligned to the human reference genome sequence (UCSC hg19) using BWA ^28^(version 0.5.9). BAM files were firstly processed by Picard (version 1.54, http://broadinstitute.github.io/picard) to sort, merge and mark duplications, and then were managed by Genome Analysis Toolkit^29^ (GATK, version 3.4) to recalibrate bases, call variants in HaplotypeCaller mode and recalibrate variant quality scores. Only variants labeled as “PASS” by GATK were kept.

### Imputation of MHC alleles

Using variants called by GATK, we imputed four-digit classical HLA alleles for HLA-A, HLA-B, HLA-C, HLA-DRB1, HLA-DQB1, HLA-DPB1, HLA-DPA1 and HLA-DQA1 with Beagle^30^ (version 4.1). The Han-MHC reference panel^31^ (total number of individuals 10,689) was used for imputation. For each sample and each gene, MHC alleles with the highest genotype probability (estimated by Beagle) were selected. SNPs corresponding to MHC alleles were obtained by aligning exon sequences from IMGT/HLA database^2^ to the human reference genome sequence (UCSC hg19) using BWA (version 0.5.9). Amino acid polymorphisms corresponding to MHC alleles were obtained by annotation of SNPs using ANNOVAR^32^ (version 2017Jul16).

### QTL mapping

The genetic variations at single nucleotide, amino acid, and four-digit classical MHC haplotypes were coded as allelic dosage counting on the number of reference allele/s (0, 1, 2). We used the MatrixEQTL R package^33^ for QTL mapping of TRBV and IGHV genes usage with genetic variations in the MHC region. We applied QTL mapping at three levels of genetic variations. SNPs, polymorphic amino acids and four-digit MHC alleles with a MAF (minor allele frequency) < 0.05 were removed. A threshold of 5% FDR was used to control for the testing burdens of multiple variations at each genetic level.

### Conditional analysis of associations between TRBV genes usage and MHC amino acid variations

The Conditional analysis was performed individually for each TRBV gene usage using forward stepwise linear regression. We established the optimal threshold to perform the stepwise conditional regression on amino acid variations with FDR < 0.05 from the QTL mapping and we considered an expanded model that included the dosage variable for target amino acid variant and the most significant amino acid variant (top amino acid variant) as a covariate. If the target amino acid variations had a conditional *P* value < 0.05, we considered it as an independent signal (second amino acid variant). The multiple linear regression model then expanded to include the top amino acid variant and the second amino acid variant as covariates and test the association of the remaining amino acid variants. Regression stopped when the conditional *P* value of the target amino acid variant was greater than 0.05. We also carried out this type of analysis for the SNPs.

### Estimating the fraction of variation for each TRBV gene usage that is explained by genetic variation in the MHC locus

We fitted a linear regression or a multiple linear regression model (when appropriate) with the target TRBV gene usage as a dependent variable and the significant amino acid variants identified from QTL mapping and conditional analyses as independent variables using *lm* function in R. The proportion of variability explained by the corresponding independent amino acid variant was the adjusted R-square derived from the linear regression model. The statistical significance of the overall model was tested using F-tests. These analyses were also performed at the level of SNPs.

### Structure analysis

Our QTL mapping results of total of 96 MHC amino acid variations were marked onto protein structures of pMHC complexes. Structures of proteins were downloaded from the RCSB PDB (PDB ID: 2BNQ^34^, PDB ID: 4G9F^35^, PDB ID: 1EFX^36^, PDB ID: 4OZF^37^, PDB ID: 1J8H^38^) and were plotted with the UCSF Chimera package^39^. We next collected all structures of TCRs bound to pMHCs and all contacting information between corresponding chains of HLA-A, HLA-B, HLA-DRB1, HLA-DQA1 or HLA-DQB1 and any of the TCR β chains or the presented peptides in the IMGT database^2^ and calculated the frequency of the presence of contacting amino acid position among the analyzed TCR-pMHC complex. The identified 96 amino acid variants were then directly compared with TCR β and peptide contacting MHC positions.

## Acknowledgments

This research was supported by the National Natural Science Foundation of China (81673181 and 31700794) and the Shenzhen Municipal Government of China (JCYJ20170817145404433 and JCYJ20170817145428361). The funders had no role in the study design, data collection, and analysis, the decision to publish or the preparation of the manuscript. We also want to thank the donors for providing UCB samples, colleagues of China National GeneBank, BGI-Shenzhen, for their help in producing the solid data, and Yafeng Zhu and Xiaowei Jiang for their help in revising the manuscript.

## Author Contribution

Xiu Qiu and Xiao Liu conceived and provided overall guidance. Lei Tian redesigned the analysis. Kai Gao, Lingyan Chen and Yuanwei Zhang carried out the analysis. Liya Lin, Jinghua Wu, Yashu Kuang, and Jinhua Lu carried out the experiments. Ziyun Wan and Yi Zhao performed the data curation. Kai Gao and Lei Tian wrote the manuscript. Xiuqing Zhang supervised the implementation of the project.

## Conflicts of interest

The authors declare no conflict of interest.

---

**Supplementary Figure 1 The frequencies of TRBV and IGHV genes.** Log2-transformed frequencies of TRBV **(A)** and IGHV **(B)**. 0.01 and 0.00001 pseudo-usage was added to avoid zeroes for TRBV and IGHV, respectively. Rows and columns were clustered using hierarchical clustering.

**Supplementary Figure 2 Independent associations between other TRBV genes and amino acid variations in MHC alleles.** Heat map representing color-coded correlation matrix of the MHC amino acids that are significantly associated with the frequencies of TRBV13 **(A)** TRBV7-9 **(B)**, TRBV10-3 **(C)**, TRBV7-6 **(D)**, and TRBV9**(E)** (FDR ≤ 0.05), and their frequencies influence by the independent amino acid variations. **(F)** The frequencies of TRBV30 gene has a single association with DQB1 residue 55.

**Supplementary table 1. TRBV gene usage matrix.**

**Supplementary Table 2. IGHV gene usage matrix.**

**Supplementary Table 3. All imputed alleles of eight MHC genes of 201 samples.**

**Supplementary Table 4. Results of QTL analysis between MHC alleles and TRBV gene usage.**

**Supplementary Table 5. Results of QTL analysis between MHC alleles and IGHV gene usage.**

**Supplementary Table 6. Results of QTL analysis between MHC SNPs and TRBV gene usage.**

**Supplementary Table 7. Results of QTL analysis between MHC SNPs and IGHV gene usage.**

**Supplementary Table 8. Results of QTL analysis between MHC amino acids and TRBV gene usage.**

**Supplementary Table 9. Results of QTL analysis between MHC amino acids and IGHV gene usage.**

**Supplementary Table 10. A list of PDB accession codes, MHC alleles and TCR V**β**-gene used in the analysis of TCR-pMHC complexes.**

