## Supplementary figures and images for "Germline-encoded TCR-MHC contacts promote TCR V gene bias in umbilical cord blood T cell repertoire"

### Supplemental Figure 1

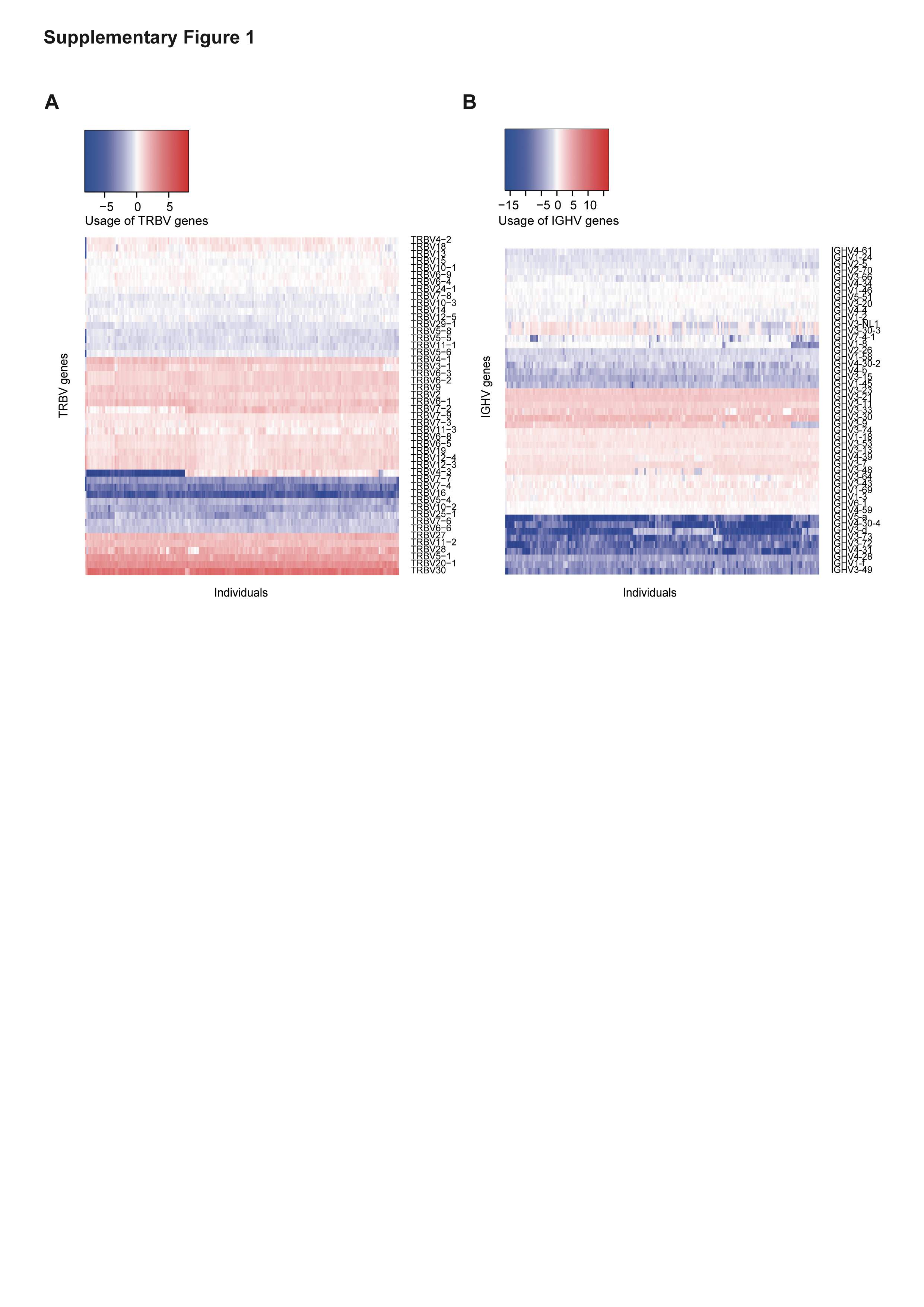

### Supplemental Figure 2

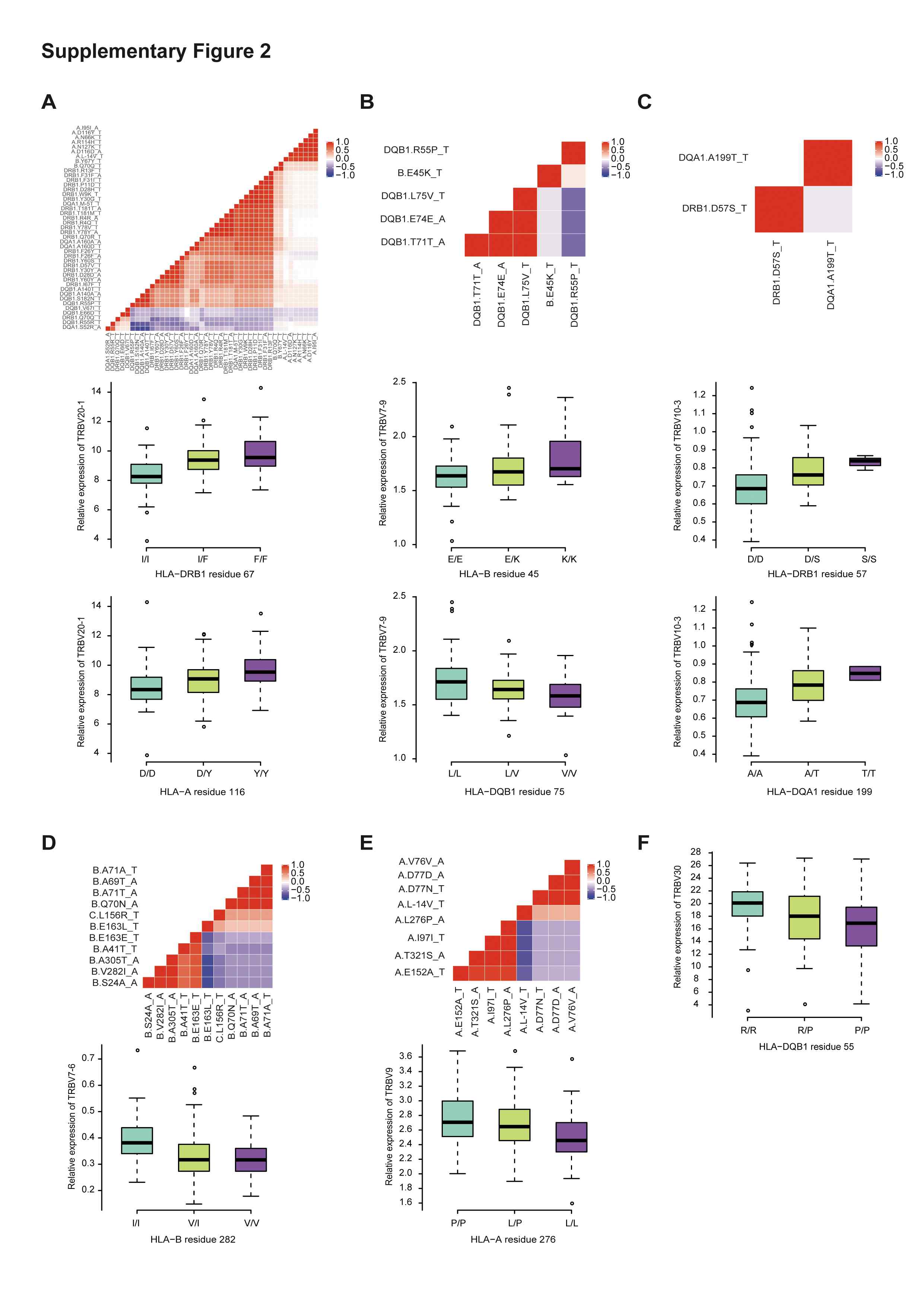
